# Decoding Predictions and Violations of Object Position and Category in Time-resolved EEG

**DOI:** 10.1101/2020.04.08.032888

**Authors:** Christopher J. Whyte, Amanda K. Robinson, Tijl Grootswagers, Hinze Hogendoorn, Thomas A. Carlson

## Abstract

Classic models of predictive coding propose that sensory systems use information retained from prior experience to predict current sensory input. Any mismatch between predicted and current input (prediction error) is then fed forward up the hierarchy leading to a revision of the prediction. We tested this hypothesis in the domain of object vision using a combination of multivariate pattern analysis and time-resolved electroencephalography. We presented participants with sequences of images that stepped around fixation in a predictable order. On the majority of presentations, the images conformed to a consistent pattern of position order and object category order, however, on a subset of presentations the last image in the sequence violated the established pattern by either violating the predicted category or position of the object. Contrary to classic predictive coding when decoding position and category we found no differences in decoding accuracy between predictable and violation conditions. However, consistent with recent extensions of predictive coding, exploratory analyses showed that a greater proportion of predictions was made to the forthcoming position in the sequence than to either the previous position or the position behind the previous position suggesting that the visual system actively anticipates future input as opposed to just inferring current input.

## Introduction

The human brain processes the position and category of objects within the visual field seemingly without effort. The process of recognising objects, although not apparent via introspection, underpins all of our interactions with the world. Even simple tasks such as making a cup of coffee rely on our ability to rapidly categorise and locate objects within the visual field. The temporal evolution of object recognition has been well characterised experimentally (Carlson, Tovar, Alink & Kriegeskorte, 2013; Cichy, Pantazis, & Oliva, 2014; Grootswagers, Robinson & Carlson, 2019; Grootswagers, Robinson, Shatek & Carlson, 2019; Robinson, Grootswagers & Carlson, 2019), however, the computational architecture underlying this process is a matter of ongoing investigation. A possible clue comes from the highly predictable structure of the visual environment. Objects tend to move along predictable trajectories giving rise to eye movement strategies such as smooth pursuit (Barnes, 2008). And, contextual knowledge of a scene greatly constrains the category of objects that are likely be to present (Bar, 2004). Given the exorbitant metabolic demands of neural processing (Stone, 2018), and the importance of prospective computation for survival (Hopfield, 1994), it would be surprising if the brain did not exploit the inherent redundancy in visual input (resulting from the structured nature of the environment) in the service of perception. In fact, for *any* system responding to an input signal that retains information from the input (i.e., has a non-zero form of memory), the retention of non-predictive information is formally equivalent to energetic inefficiency (Still, Sivak, Bell & Crooks, 2012). The question is, therefore, not whether brains predict but how.

The prospective goal of perception lies at the heart of a family of models in computational neuroscience collectively referred to as predictive processing models (Clark, 2016). Predictive processing models, which includes both predictive coding (Bastos et al, 2012; Friston, 2005; Friston & Kiebel, 2009; Rao & Ballard, 1999) and active inference (Friston et al., 2017; Parr, Da Costa & Friston, 2020), have shown great promise in accounting for a wide range of visual phenomena from extra-classical receptive field effects (Rao & Ballard, 1999) and repetition suppression (Auksztulewicz & Friston, 2016), to selective attention (Feldman & Friston, 2010; Mirza, Adams, Friston & Parr, 2019), and even visual awareness (Parr, Corcoran, Friston & Hohwy, 2019; Whyte & Smith, 2020). Of the predictive processing models, by far the most popular is the classic predictive coding model proposed by Rao and Ballard (1999) and later built upon by Friston (2005). Unlike feed-forward neural networks, predictive coding depicts perception as a process of top-down model testing aimed at minimising the difference between an internal model of the world and sensory input. The internal model generates cascades of descending predictions that meet bottom-up signals at each level of the visual hierarchy. The mismatch between the prediction and the bottom-up signal (prediction error) is fed forward to the next level in the hierarchy leading to a revision of the prediction. (Bastos et al, 2012; Friston, 2005, 2010; Clark, 2016; Hohwy, 2013, Rao & Ballard, 1999). In line with this view, there is now considerable evidence from functional magnetic resonance imaging (fMRI) suggesting that prediction has a silencing effect on neural responses that is orthogonal to other top-down processes such as attention (Kok et al, 2011; Richter, Ekman & de Lange, 2018), and that higher levels in the cortical hierarchy send predictions to subordinate levels of the hierarchy (Summerfield et al, 2006).

Also in line with predictive processing models, a body of research conducted in magnetoencephalography and electroencephalography (M/EEG) has shown that expectation also has substantial effects in the temporal domain. When stimuli are expected, stimulus features such as orientation can be decoded even before stimulus onset (Kok, Mostert & De Lange, 2017). Decoding of object position in apparent motion paradigms has a latency advantage when the target stimulus moves along a predictable trajectory (Hogendoorn & Burkitt, 2018), and the violation of the orientation and identity of faces has a dissociable mismatch ERP effect across the dorsal and ventral streams (Robinson et al., 2018).

Here we used time-resolved multivariate pattern analysis and EEG (MVPA; Carlson et al, 2013; Cichy et al, 2014; Grootswagers, Wardle & Carlson, 2017) to investigate the temporal effects of prediction and prediction error at different levels of the visual hierarchy. We presented participants with sequences of images that stepped around fixation in a predictable order. On the majority of sequences, the images conformed to the pattern of position and category order, however, on a subset of the sequences the last image in the sequence violated the established pattern by violating either the predicted category (high level) or the predicted position (low level) of the object.

Based upon the classic formulation of predictive coding (Friston, 2005; Rao & Ballard, 1999), and the structure of visual hierarchy (Felleman & Van Essen, 1991), we generated four related hypotheses (see Figure 1). First, we expected that due to prediction error signals, there would be above chance decoding between predicted stimuli, and stimuli that have violated a prediction, for both violations of position and category. Second, since the generation of prediction error is hypothesised to alter the content of representations (c.f., King, Schurger, Naccache & Dehaene, 2014) we expected representations to be less separable when predictions were violated leading to reduced decodability. Third, given the relative independence of the dorsal and ventral streams (Ungerleider & Haxby, 1994) we expected that category violation, which relies on processing within the ventral stream, would not interact with position representation, which relies on processing within the dorsal stream, and vice versa. Several aspects of this study including hypotheses, design, and analysis choices were pre-registered (https://osf.io/hkedz/).

**Figure 1.**
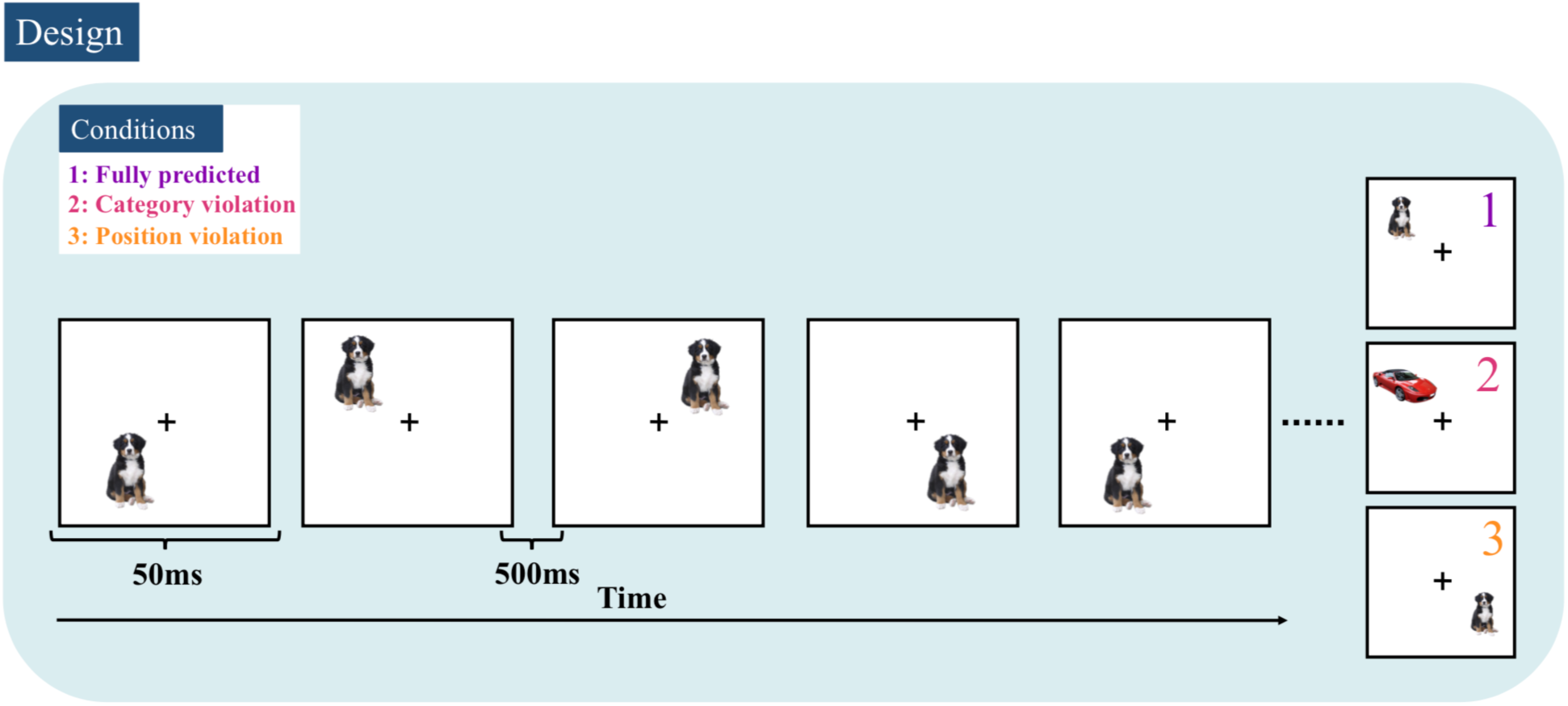
Experimental design. Stimuli were presented for 50ms with 500ms ISI in sequence around four possible locations that were equidistant from the fixation cross and either repeated or alternated the category of the stimulus (i.e., dog or car). The last image in the sequence either conformed to the pattern (1: fully predicted - purple), or violated the established pattern by violating either the the predicted category (2: category violation - pink), or the predicted position (3: position violation - yellow). This example shows a clockwise repeating object sequence.

## Methods

### Stimuli and procedure

We recruited 34 adult participants (21 female) aged between 18 – 27 years old (average 20.15) from the University of Sydney in exchange for course credit or payment. Participants viewed sequences of dog and car images (obtained from the free image site www.pngimage.com) that appeared in four different positions. Stimuli were presented in sequences of 6 to 10 images that stepped around fixation in a predictable order (50% clockwise, 50% counter clockwise). The stimulus subtended 3×3 degrees of visual angle and was presented 4 degrees from fixation. Each stimulus was presented for 50ms with a 500ms inter-stimulus interval. There were two types of predictable sequences: repeating object sequences (50%) and alternating object sequences (50%). During the repeating sequences, the same stimulus (dog or car) was presented throughout the sequence. In the alternating sequences the category of stimulus alternated on each successive presentation in the sequence (e.g., dog, car, dog, car…). There were 490 sequences in total. On the majority of sequences (256 out of 448 non target sequences), all stimuli conformed to the pattern of position order (clockwise/counter-clockwise) and object order (repeating/alternating). For the remaining sequences, the last image in the sequence violated the established pattern by either violating the predicted category of the object (*category violation*; e.g., dog-dog-dog-***car*** or dog-car-dog-***dog***; 96 sequences) or the predicted location (*position violation*; 96 sequences). See Figure 1. Importantly, for position violation sequences the position of the last stimulus was a reversal of the established movement (e.g., positions 4-1-2-3-***2*** or 1-4-3-2-***3***. This ensured that for all conditions, the previous item in the sequence could not be a confound in the decoding analysis.

Participants were required to monitor the sequence for inverted stimuli which appeared 8.57% of the time (42 sequences). They were instructed to fixate on the cross in the centre of the monitor, and not to move their eyes. The inversion task kept them alert and attentive without making the predictability of the stimulus task relevant. With the exception of the inversion of the target stimulus, target sequences were identical to predictable non-target sequences. Target sequences (i.e., sequences with inverted stimuli) were excluded from analysis.

Participants completed 7 blocks each consisting of 70 sequences. Between each block we presented a ‘pattern localiser’ consisting of a rapid stream of 120 dog and car images yielding a total of 840 additional presentations (i.e., 12 repeats of each dog and car image at each of the 5 locations). Each image was presented with 50ms ISI and 100ms SOA either centrally (at fixation) or at one of the four experimental locations. The location of the stimulus was shuffled such that there was no statistical regularity in the sequence. The pattern localiser served as an independent source of training data for the decoding analysis.

### EEG Recordings and Pre-processing

Continuous EEG data was recorded with a BrainVision ActiChamp system with a digitised sampling rate of 1000Hz. The 64 electrode system was arranged according to the 10-10 placement system all referenced to Cz. Pre-processing was conducted in MATLAB using the EEGLAB toolbox (Delorme & Makeig 2004). The data were filtered with a high pass filter of 0.1Hz and a low pass filter of 100Hz and down-sampled to 250Hz. Epochs were created between -200 to 1000ms relative to the onset of the final image in the sequence (448 epochs).

### Decoding Analysis

We employed an MVPA decoding pipeline to all EEG channels following the recommendations of Grootswagers, Wardle and Carlson (2017) using the CoSMoMVPA toolbox (Oosterhof, Connolly & Haxby, 2016). All decoding was performed within subject using a linear discriminant analysis (LDA) classifier. Statistical analysis was performed at the group level averaging across individual decoding accuracies. To explore the emergence of prediction error signals we compared neural responses of violation trials with neural responses of predictable trials. For violation decoding we used a leave - one block - out cross validation scheme. There were two separate analyses: predictable versus object violation and predictable versus position violation. There were far more predictable sequences, so to ensure balanced data in the test set for every unpredictable trial, we selected a predictable trial that was matched for repeating/alternating sequence, clockwise/counter clockwise sequence, category and position. For *position* decoding we used a cross decoding scheme by training the classifier on data from the pattern localiser using the four peripheral positions and testing the classifier on data from the experimental sequences. For category decoding we again used a cross decoding scheme training on data from the pattern localiser and testing on data from the response sequences. However, for category decoding we trained the classifier on stimuli presented at all 5 locations of the pattern localiser (4 peripheral positions and central) to get a better estimate of position-invariant image category information.

In total we decoded 8 contrasts; *violation* split by category and position (contrasts 1-2); *position* (i.e., location 1-4) split by fully predicted, position violation, and category violation conditions (contrasts 3-5) and *category* (i.e., dog vs car) fully predicted, position violation, and category violation conditions (contrasts 6-8).

### Exploratory Analysis of Classification Errors

At the time point of peak decoding for position we found insufficient evidence to determine if there was more position information in the neural signal for predictable compared with position violation trials (hypothesis 2; Figure 2). In order to investigate this hypothesis further, we examined the predictions made by the classifier. The classifier extracted neural patterns of activation specific to each of the four experimental positions and used these to predict the position of the stimulus on each experimental trial. If there was no position information about the stimulus (e.g., prior to its appearance), the classifier would be expected to predict each of the four locations equally often. If there was position information in the neural signal (e.g., during retinotopic processing of the stimulus), the classifier would be expected to predict the correct position. However, classification is rarely perfect, and investigating the errors made by the classifier can give insight into other information in the neural signal. For example, in an apparent motion paradigm Blom, Feuerriegel, Johnson, Bode and Hogendoorn (2020) trained a classifier to decode between stimuli presented at the two locations on either side of the target stimulus and then tested the classifier on the location of the target stimulus. For the first ∼70ms the majority of the predictions made by the classifier were made to the location behind the location of the target but after ∼70ms the majority of predictions were made to the location following the target showing that there was anticipatory information in the EEG signal.

**Figure 2.**
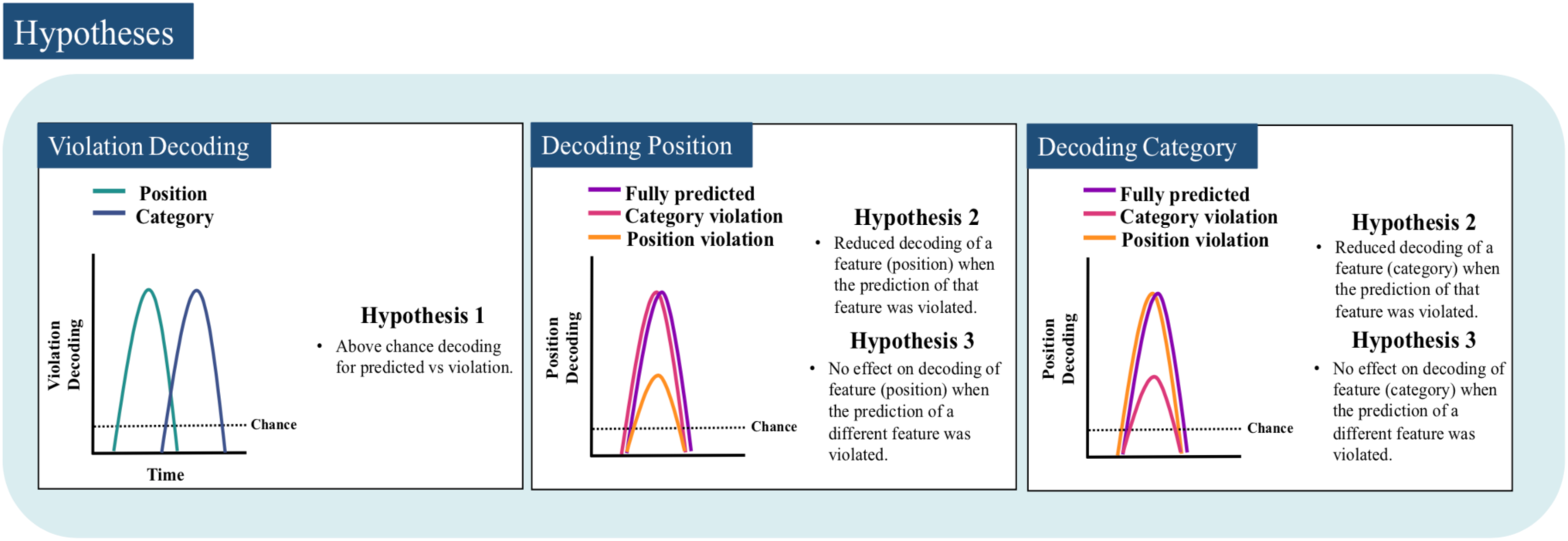
Hypotheses for each condition. Hypothesis 1 – the increase in prediction error on violation trials was expected to lead to above chance decoding between prediction and violation trials for both position and category. Category violation decoding is shifted rightward in the figure because category is extracted at a higher point of the hierarchy than position so the appearance of prediction errors related to category processing is expected to occur later in time. Hypothesis 2 – given that prediction error leads to a revision of the content of representations we expected representations to be less separable when predictions were violated thereby lowering decoding accuracy. Hypothesis 3 – given the relative functional independence of the dorsal and ventral streams we expected that the violation of a feature that is not the target of decoding would have no effect on decoding accuracy.

We examined the average proportion of predictions made by the classifier to each position. We then sorted the predictions made by the classifier to each of the four locations relative to the expected position. Assuming there was only information about the current stimulus in the EEG signal there should have been an equal proportion of predictions made to each of the three incorrect positions. If, however, the EEG signal also contained information about one of the incorrect positions the classifier might predict one of the incorrect positions more often than the others. For example, when the stimulus was presented in an unexpected position, as was the case on violation trials, predictive information in the signal might have increased the proportion of predictions made by the classifier to the predicted position. To statistically evaluate the evidence for differences in the proportion of predictions made by classifier we took the average proportion of predictions made over a 20ms time-window (86-106ms) centred on the point of peak decoding accuracy (96ms) and used Bayes factors (described below) to evaluate the strength of evidence.

It is important to note that unlike the analyses listed above, this analysis was not planned a priori and is therefore considered exploratory.

### Statistical Inference

To calculate the evidence for the null and alternative hypotheses we used JZS Bayes Factors (Rouder et al., 2009). To determine the evidence for the alternative hypothesis of above chance decoding we employed a Cauchy prior with the scale factor set to 0.707, while the prior for the null hypothesis was a point at chance, 0.25 for position decoding and 0.5 for all other decoding tests (Morey & Rouder, 2011). To determine the evidence for a non-zero difference between decoding accuracies, we used a uniform prior with a point null set to zero. This same procedure was also used in the exploratory analysis described above. Using these distributions, we computed Bayes factors (Dienes, 2011; Jeffreys, 1998; Rouder, Speckman, Sun, Morey, & Iverson, 2009; Wagenmakers, 2007) which, being a ratio of marginal likelihoods, measures the evidence for the alternative hypothesis relative to the null. For the purpose of plotting the results we thresholded the Bayes factors at BF > 1/3 but < 3 as inconclusive evidence either way, BF > 6 for modest evidence for the alternative hypothesis, and BF > 10 for strong evidence for the alternative hypotheses. Because point nulls are biased to the alternative hypothesis as sample size becomes larger (Morey & Rouder, 2011), we took BF < 1/3 as strong evidence in favour of the null.

## Results

### Behaviour

Participants performed an orthogonal task to detect inverted stimuli. After inspecting the behavioural responses, we excluded one participant who did not respond to any of the targets. After exclusion, mean accuracy was 91.9% (SD = 10.13). We then excluded another two participants from further analysis whose accuracy was lower than 80%, leaving a total of 31 participants whose data was used in the decoding analysis. We used the inclusion criterion of above 80% accuracy because of the extreme simplicity of the task.

### Violation decoding

We expected the neural signal to contain information about whether a stimulus violated the expected pattern for both position and object (Hypothesis 1). In support of hypothesis 1 we observed strong evidence that stimuli were processed differently if they appeared in unexpected positions (position *violation*) relative to expected positions (Figure 3; green line). Position *violation* decoding was above chance (BF > 10) 244ms after stimulus onset with two peaks in accuracy at 258ms and 812ms. Each peak coincided with an increase in evidence for above chance decoding (BF > 10). In contrast, partially contrary to hypothesis 1, we did not observe a difference in processing of stimuli when the category violated the established pattern (*category violation)* relative to expected category (Figure 3; blue line). In decoding of violation for category we found strong evidence for the null hypothesis across the trial (BF < 1/3). Unsurprisingly, when comparing accuracy between the two conditions we found strong evidence (BF > 10) for a difference in decoding accuracy between the two conditions that coincided with peaks in decoding accuracy for *position* violation decoding.

**Figure 3.**
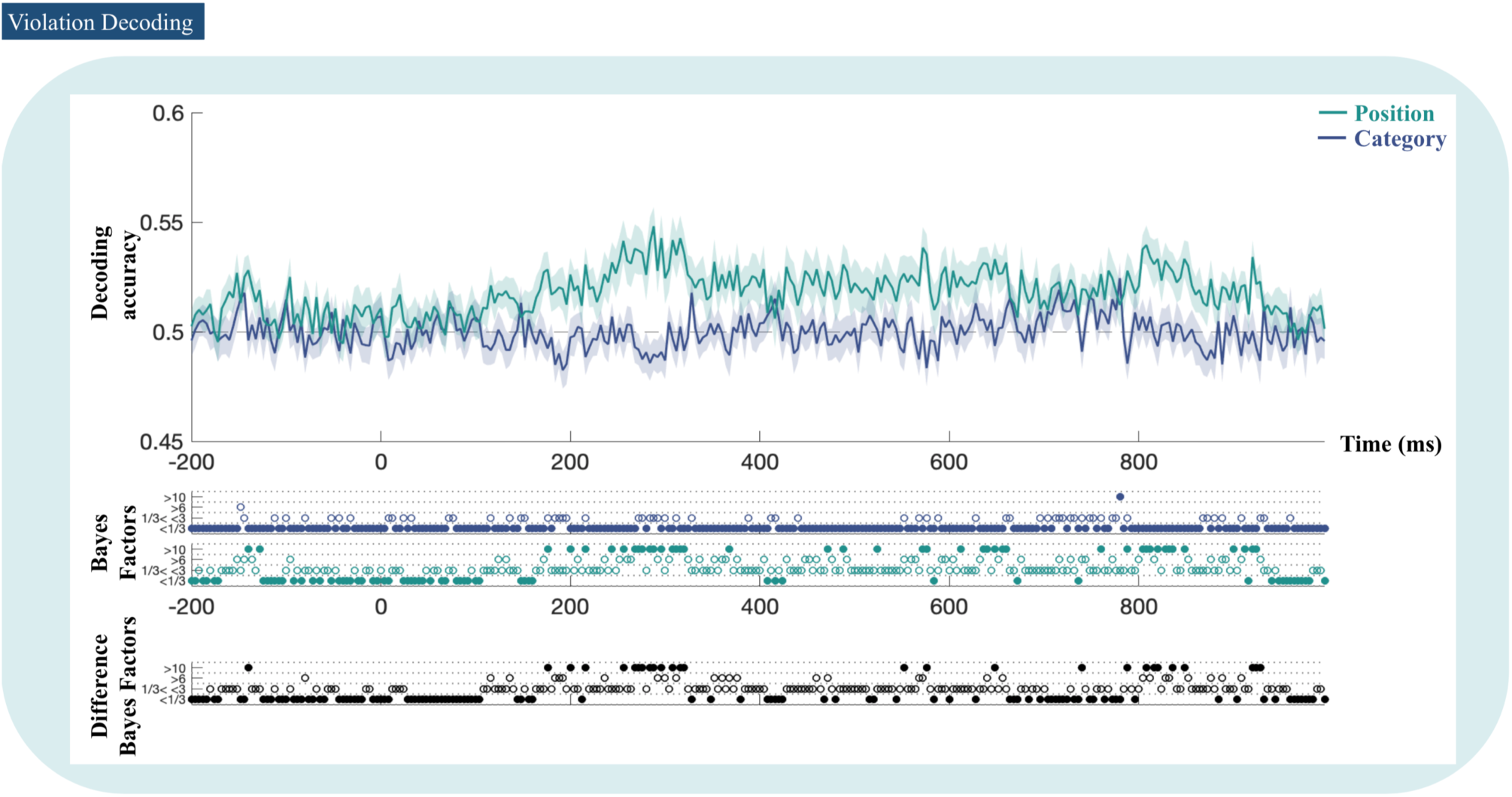
Violation decoding. Mean decoding accuracy for predictable versus position violation (green) and predictable versus category violation (blue). Coloured dotes below the plot indicate the thresholded Bayes factors (BF) for category and position. For BF > 10 and BF < 1/3 which indicate strong evidence for either the alternative or null hypothesis are shown represented by filled in circles. BF > 6 and BF <3 which indicate modest or inconclusive evidence either way are represented by open circles. Black dots indicate the thresholded BF for the difference in decoding accuracy between violation types.

### Decoding position

We expected position-related information to be present in the neural signal from an early stage of processing, and for the amount of information to differ for predictable positions compared to position violations (Hypothesis 2). We found strong evidence (BF > 10) for above chance decoding of position (chance = 25%) for *fully predicted, category violation*, and *position violation* from 72ms post stimulus onset with a peak at 96ms (Figure 4a). Interestingly, suggestive of hypothesis 2 concerning the disruptive effect of prediction error, peak decoding accuracy for *position violation* was lower than *fully predicted* and *category violation* and was less sustained. However, during the time period where there was a visible difference between decoding accuracies (∼85 to 120ms) we found inconclusive evidence for the null and alternative hypothesis (1/3 > BF < 3). To investigate this in more detail we examined the proportion of predictions made by the classifier for each location (Figure 5). If hypothesis 2 was correct, we would have expected the neural signal to contain information about the predicted position on both violation and predicted trials, indexed by higher numbers of classifier errors for the expected position than other incorrect positions on position violation trials. However, this is not what we found. Intriguingly, at the point of peak decoding accuracy (∼96ms), classification error analysis showed a higher number of predictions for the position that followed the expected position (i.e., expected + 1, the next position in the sequence) across all three conditions making up ∼25% of classifier output (green line in Figure 5). To evaluate this statistically we examined the differences between the average proportion of classifier predictions made to each position averaged over a 20ms time-window (86-106ms) centred on the point of peak decoding accuracy (96ms). For the *fully predicted* and *category violation* conditions we observed strong evidence BF > 10 that a greater proportion of predictions was made to the next position in the sequence (expected + 1) in comparison to both the previous position (expected – 1) and the position behind the previous position (expected - 2). Similarly, for the *position violation* condition we found strong evidence (BF > 10) that a greater proportion of predictions was made to the next position in the sequence (expected + 1) than to the expected position, and modest evidence (BF > 3) that a greater proportion of predictions was made to the next position than to the previous position (expected – 1). We consider the interpretation and significance of this result in the discussion section. Finally, it is worth highlighting that we could not evaluate hypothesis 3 – which predicted that there would be no difference in decoding accuracy when the violated feature was not the target of decoding – as it relied on violation having a disruptive effect as predicted by hypothesis 2.

**Figure 4.**
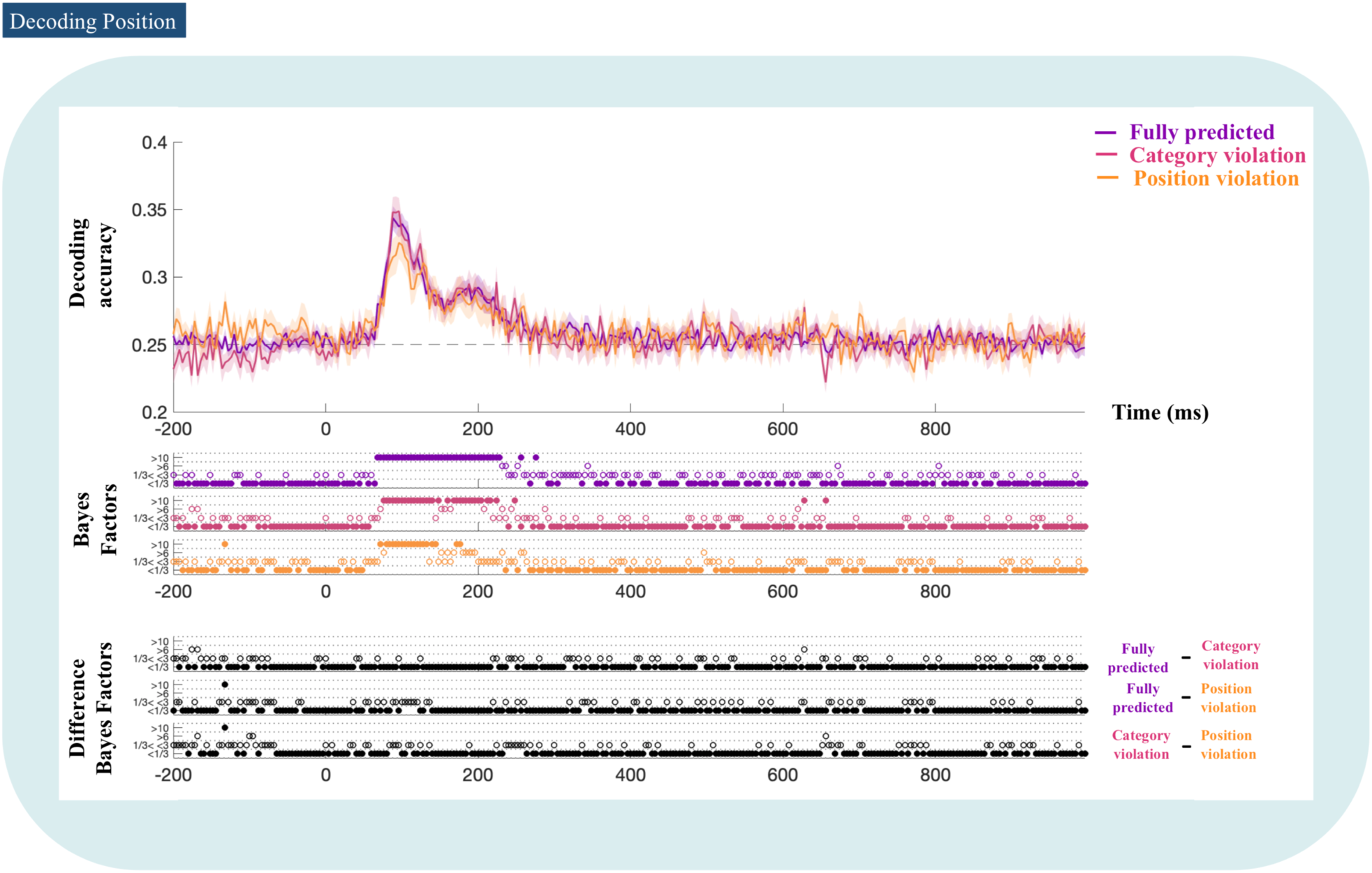
Position decoding (chance = 25%) for fully predicted, category violation and position violation conditions. Coloured dots below each plot indicate the thresholded Bayes factors for each time point. Black dots indicate the thresholded Bayes factors for the difference in decoding accuracy between conditions.

**Figure 5.**
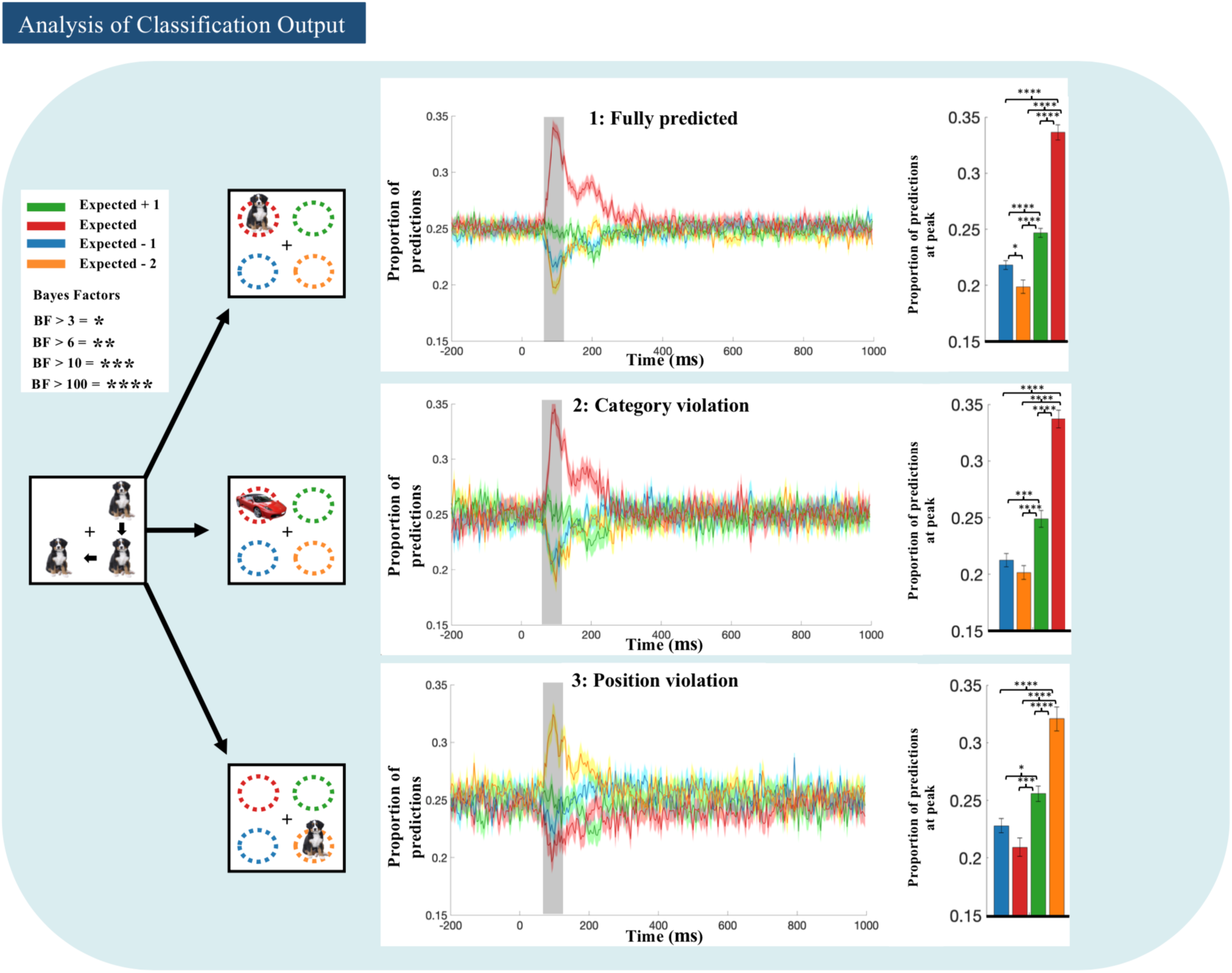
Analysis of classifier output for position decoding across conditions. Left portion of the figure shows an example of the position of the stimulus for each condition in relation to what was expected given the pattern of the preceding sequence. Colours on plots correspond to the stimulus positions. Middle graphs show the proportion of predictions made by the classifier to each position. Across conditions the actual position of the stimulus had the highest proportion of classifier predictions (∼32-34%). The right bar plots show the proportion of classifier predictions made to each position at the peak of decoding (shaded portion of graph). Asterisks* above the plots indicate the Bayes factors for the differences in proportion of predictions for each position. Contrary to hypothesis 2, there was a greater proportion of predictions made to the expected + 1 (next) location than either of the other two incorrect locations across conditions. Crucially, in the position violation condition there was exceptionally strong evidence (BF >100) that there was a greater proportion of predictions made to the expected + 1 location than the expected location suggesting the presence of anticipatory information in the EEG signal.

### Decoding Object Category

To assess object category information, we decoded car versus dog for the three predictability conditions (see Figure 6). For the *fully predicted* condition there were 6 time points between 228-260ms that showed modest (BF > 3) to strong evidence (BF > 10) for above chance decoding of *category*. Similarly, for the *category violation* condition there were 3 time points between 226-276ms that showed modest (BF > 3) to strong evidence (BF > 10) for above chance decoding, and for the *position violation* condition, we found strong evidence for the null hypothesis throughout the trial (BF < 1/3) with only a few time-points transiently showing evidence for above chance decoding. In terms of differences in decoding accuracy between conditions, with the exception of a few sparsely distributed and isolated time points, we found strong evidence for the null hypothesis that there was no difference in decoding accuracy from stimulus onset until the end of the epoch (BF < 1/3). These results stand in opposition to hypothesis 2 which forecast that violations of predictions would decrease decoding accuracy if the violation was the target of decoding. Again, we could not evaluate hypothesis 3 as it relied on violations having a disruptive effect on decoding accuracy as predicted by hypothesis 2.

**Figure 6.**
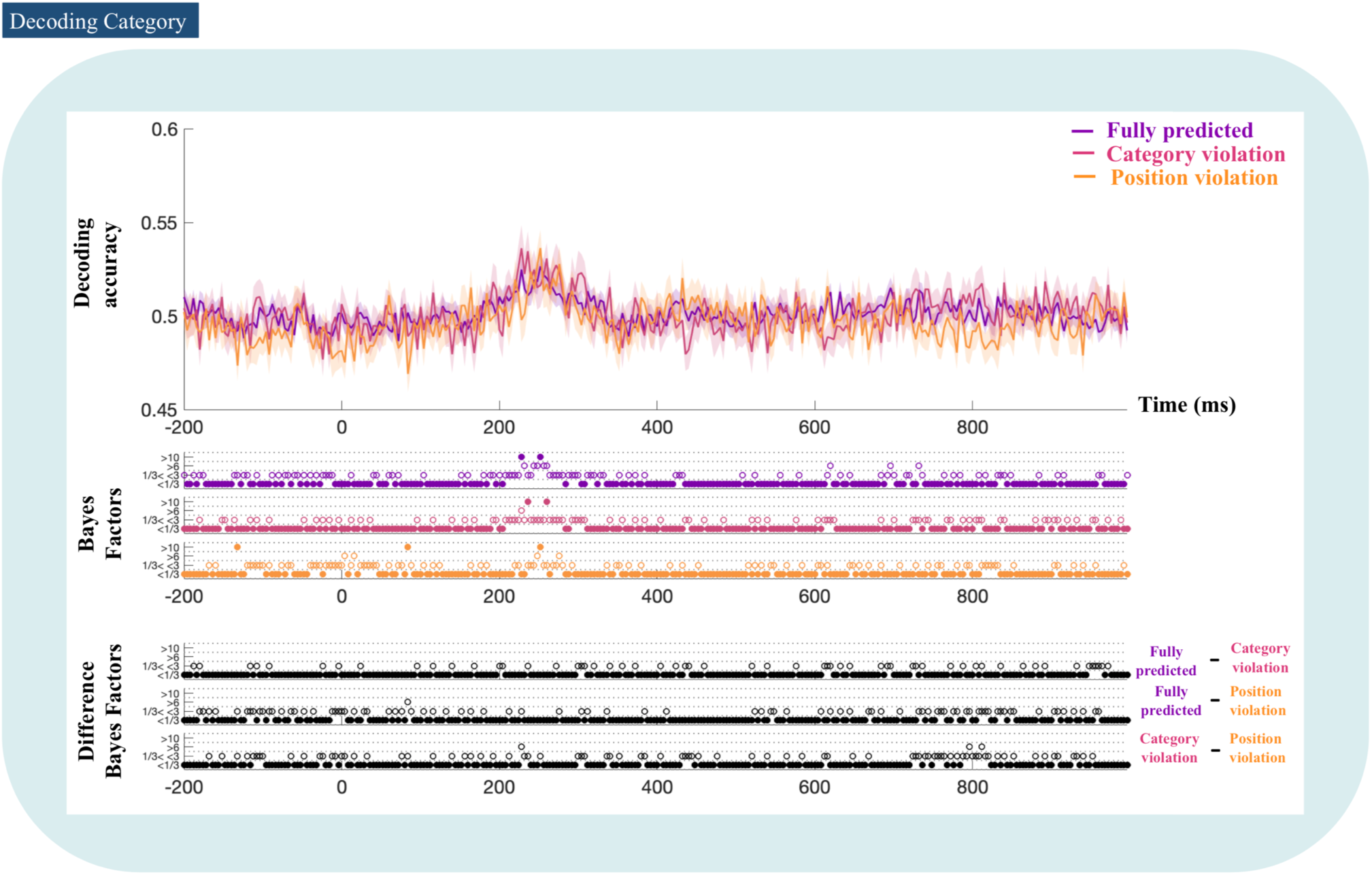
Category decoding (chance = 50%) for fully predicted, category violation and position violation conditions. Coloured dots below each plot indicate the thresholded Bayes factors for each time point. Black dots indicate the thresholded Bayes factors for the difference in decoding accuracy between conditions.

## Discussion

The aim of this study was to test a set of four hypotheses based on the classic model of predictive coding (Friston, 2005; Rao & Ballard, 1999), and the structure of the visual hierarchy (Felleman & Van Essen, 1991), in the domain of basic object vision using time-resolved EEG. According to the classic model of predictive coding sensory systems use information from prior experience to predict current incoming sensory input, or more concisely, they are trying to predict the present (Clark, 2016). Overall, our results were largely contrary to this idea. The remainder of this paper will discuss the consequence of our results for each hypothesis and propose an explanation of the findings in terms of generalised predictive coding (Friston & Kiebel, 2009; Friston, Stephan, Li & Daunizeau, 2010) and the temporal realignment hypothesis (Hogendoorn & Burkitt, 2019).

According to generalised predictive coding (Friston & Kiebel, 2009; Friston, Stephan, Li & Daunizeau, 2010), instead of just aiming to predict current input, predictions are cast in generalised coordinates of motion meaning that predictive signals also represent the velocity, acceleration, and other higher order derivatives of the predicted input allowing sensory systems to extrapolate. For an accessible introduction to the idea of generalised coordinates of motion see Susskind & Hrabovsky, 2014, and for its role in predictive coding see Buckley, Kim, McGregor & Seth, 2017. Similarly, the temporal realignment hypothesis (Hogendoorn & Burkitt, 2019) proposes that the brain overcomes the transmission delays inherent to the visual system by having both predictions and prediction errors extrapolate forward in time. The key feature of both of these models is that they posit the existence of temporal predictive signals that carry information about what *will* happen and not just what *is* happening.

For *prediction vs violation* of position, we observed strong evidence for above chance decoding which peaked at 258ms. Prima facie, this result seems in line with hypothesis 1 which predicted that the presence of prediction error signals on the violation trials would lead to above chance *violation* decoding. However, decoding was far too late to plausibly reflect an error message which by hypothesis would occur at a similar time-point as peak decoding accuracy for position. In fact, the peak in decoding accuracy for *violation* of position occurred ∼150ms later than peak decoding accuracy for position. As such, the time course of the response is more likely due to an orienting of attention (Carlson, Hogendoorn & Verstraten, 2006; Eimer, 2000).) to the unpredicted position. In favour of this interpretation, for *violation* of category we found strong evidence for the null hypothesis of no above chance decoding. If violations of predictions generated error signals that were large enough to be detected at the level of the scalp we would have expected to see above chance decoding of *violation* for category as well as position.

Assuming the interpretation put forward above is on track, the lack of a decodable error signal suggests that prediction and prediction error may have a subtler effect than we initially hypothesised. Indeed, considering that the stimulus moved around the screen and did not stay within a consistent set of receptive fields, the short-term changes in synaptic plasticity that are thought to underlie the generation of error related ERPs (Auksztulewicz & Friston, 2016; Garrido, Kilner, Stephan & Friston, 2009; Stefanics, Kremláček & Czigler, 2014) would have been reduced, and as such, the changes in voltage that characterise violation effects in ERP would have been less pronounced.

With that said, it is still plausible that the presence of predictive signals could have caused the classifier to make more (erroneous) predictions to the expected position on violation trials. Although again, this is not what we found. When decoding position, all three conditions - *fully predicted, category violation*, and *position violation* - displayed above chance decoding accuracy with a peak at 96ms. Importantly, peak decoding accuracy for *position violation* seemed lower than *fully predicted* and *category violation* suggesting that prediction error may have had a disruptive effect on position information as proposed in hypothesis 2. However, we found inconclusive evidence differentiating between the null and alternative hypotheses at this time point. To investigate the effect of position violation on position coding in a different way, we inspected the classification output for each of the three conditions. If hypothesis 2 was correct we would have expected to see a greater proportion of predictions to be made to the expected location. Instead, the classifier made a greater proportion of predictions to the next position in the sequence (expected + 1) across all three conditions. Against the classic model of predictive coding this suggests that the visual system actively anticipates future input as opposed to just inferring current input. Crucially, however, this finding is predicted by both generalised predictive coding (Friston & Kiebel, 2009; Friston, Stephan, Li & Daunizeau, 2010), and the temporal realignment hypothesis (Hogendoorn & Burkitt, 2019), which propose that predictions extrapolate forward in time. Further, our results mirror those of Blom, Feuerriegel, Johnson, Bode and Hogendoorn (2020), who found that when a stimulus was a part of a predictable sequence information about of the stimulus’ next location was present in the EEG signal 70 - 90ms earlier than would be expected if the evoked response was purely stimulus driven.

We modest evidence for above chance classification of category in all three conditions. However, contrary to hypothesis 2, which forecast that violations of predictions would show lower decoding accuracy, we found strong evidence for the null hypothesis that there were no differences between conditions. The lack of effect for category violation has at least two plausible and complementary explanations. First, it may simply be that there was an effect of violation at the neuronal population level but because the cortical representation of objects is weaker in the peripheral parts of the visual field where our stimuli were presented (Levy, Hasson, Avidan, Hendler, & Malach, 2001) the differences could not be seen at the level of the scalp. Indeed, we observed clear category decoding, yet the absolute decoding accuracy was fairly low compared with previous studies using centrally presented objects (e.g. Grootswagers, Robinson & Carlson, 2019; Grootswagers, Robinson, Shatek & Carlson, 2019; Robinson, Grootswagers & Carlson, 2019). Second, like position, it may be that predictions of category are primarily anticipatory in nature and as such we should expect to see a greater proportion of classification errors made to the category of the next stimulus but not the current stimulus. Unfortunately, however, our paradigm did not allow us to interrogate this hypothesis. Since we only had two stimulus categories and our stimulus was presented in an alternating or repeating pattern, the category of the upcoming stimulus was either the same as the current category or the same as the previous category making classifier output uninformative. Still, this hypothesis will be easy to test in future work by simply increasing the number of stimulus categories and presenting the stimuli in predictable sequences at the centre of the screen where there is a stronger cortical representation of object category (as has been done in fMRI; Richter, Ekman & de Lange, 2018). Relatedly, in terms of hypothesis 3, which forecast that violations of the non-target feature would have no effect on decoding, we cannot evaluate its accuracy as it relied on violations having a disruptive effect on decoding (i.e., hypothesis 2). Unfortunately, if we are correct in arguing that the effect of prediction error is reduced the periphery, this hypothesis will be difficult to test using non-invasive techniques.

In sum, our results are largely contrary to the classic model of predictive coding (Friston, 2005; Rao & Ballard, 1999) which proposes that sensory systems use prior experience to predict the present (cf. Clark, 2016). Instead, consistent with generalised predictive coding, and the temporal realignment hypothesis, our exploratory analysis suggests that sensory signals are actively anticipate future input, at least for representations of position. This contrary finding, which was predicted by previous theoretical work, represents an important advance in how we should think about prediction in sensory systems. We look forward to future work investigating whether the anticipatory nature of prediction generalises to category representation.

## Acknowledgments

We are grateful to Soyoun Park and Christopher Makin for assistance in data collection.

